# Adapted dandelions increase seed dispersal potential when they are attacked by root herbivores

**DOI:** 10.1101/551630

**Authors:** Zoe Bont, Marc Pfander, Christelle A. M. Robert, Meret Huber, Erik H. Poelman, Ciska E. Raaijmakers, Matthias Erb

## Abstract

Plants allow their offspring to escape unfavourable local conditions through seed dispersal. Whether plants use this strategy to escape herbivores is not well understood. Here, we explore how different *Taraxacum officinale* populations modify seed dispersal in response to root herbivore attack by *Melolontha melolontha* in the field. Root herbivore attack increases seed dispersal potential through a reduction in seed weight in populations that have evolved under high root herbivore pressure, but not in populations that have evolved under low pressure. This increase in dispersal potential is associated with reduced germination, suggesting that adapted plants trade dispersal for establishment. Analysis of vegetative growth parameters suggests that increased dispersal is not the result of stress flowering. These results suggest that root herbivory selects for genotypes that increase their dispersal ability in response to herbivore attack.

## Introduction

As sessile organisms, plants are bound to grow and develop where they germinate. Local conditions are therefore important determinants of plant survival and reproductive success (Kawecki & Ebert 2004, Walter et al. 2018). Although the ability of plants to move is limited, they can influence the displacement capacity of their offspring by modifying dispersal traits (Nathan & Muller-Landau 2000, Martorell & Martínez-López 2014, Teller et al. 2014). Theory suggests that whenever local conditions are unfavorable, increased seed dispersal is advantageous, as it allows the next generation to escape from the unsuitable environment and increases their chance to colonize new habitats with better conditions (Levin et al. 2003, Ronce 2007). Major dispersal strategies of plants include dispersal by wind (anemochory), water (hydrochory) and animals (zoochory) (Poschlod et al. 2013).

Herbivores can have strong impacts on plant development and fitness and thus act as a force of natural selection (Rutter & Rausher 2004, Lau et al. 2008, Agrawal et al. 2012, Züst et al. 2012). To date, the impact of herbivores on plant dispersal remains poorly understood (de la Peña et al. 2011). Compared to aboveground herbivores, belowground herbivores are generally more limited in their mobility due to the dense nature of soil and are therefore often distributed in a patchy way (Van der Putten et al. 2001). Hence, seed dispersal may allow plants to escape belowground enemies, which may lead to selection for high dispersal genotypes by root herbivores (Van der Putten 2003). Experimental evidence for this hypothesis is currently lacking. Understanding how herbivores influence seed dispersal and establishment traits would help to elucidate their impact on plant population dynamics. In general, herbivores may affect plant dispersal strategies either by selecting for genetically fixed high dispersal genotypes or by favoring genotypes that are phenotypically plastic and increase dispersal in response to herbivore attack. Especially in stochastic environments with fluctuating herbivore pressure, inducible increase of seed dispersal distance as plastic response to herbivory may be advantageous (Jakobsson & Eriksson 2003).

Trade-offs between dispersal capacity and seed viability are frequent in nature. For distribution processes of wind-dispersed plant species, seed mass is often reported as a critical trait that influences both dispersal and seedling establishment. Lower seed mass typically increases dispersal (Meyer & Carlson 2001; Greene & Quesada 2005; Skarpaas et al. 2011), but reduces germination and early growth (Leishman & Westoby 1994; Turnbull et al. 2004). Thus, by varying seed mass, wind-dispersed plant species may trade dispersal distance for establishment or *vice versa*. Studying seed dispersal capacity together with establishment traits is, therefore, essential to understand how seed-dispersal influences plant population dynamics.

Here, we studied the ecological and evolutionary impact of root herbivory on seed dispersal and fitness in the common dandelion, *Taraxacum officinale* agg. (Asteraceae). *T. officinale* is a species complex native to Eurasia that consists of sexual, obligate outcrossing diploids and asexual, apomictic triploids. *T. officinale* produces seeds with characteristic parachutes (pappus), which enhance uplift by the formation of separated vortex rings after detachment (Cummins et al. 2018). The seeds can be dispersed efficiently by wind and allow the plant to rapidly colonize new habitats (Tackenberg et al. 2003). A recent study found that *T. officinale* exposed to aboveground herbivory produced seeds with longer pappus and thus increased dispersal ability (de la Peña & Bonte, 2014). However, as locust exposure also results in higher biomass production, it is unclear if the change in seed morphology is a direct plastic response to herbivore exposure or an indirect consequence of changes in plant growth (de la Peña & Bonte, 2014). Furthermore, whether herbivory-induced changes in seed traits are related to the plant’s evolutionary history, i.e. whether they are the result of natural selection by herbivore attack, is unclear. The larvae of *Melolontha melolontha* (Coleoptera: Scarabaeidae), also called white grubs, are the major native belowground herbivores of *T. officinale.* Although the larvae are polyphagous, they prefer to feed on *T. officinale* (Hauss 1975, Hauss & Schütte 1976). In a recent study, we showed that *T. officinale* populations that evolved under high *M. melolontha* pressure over the last decades produce higher concentrations of deterrent root secondary metabolites, suggesting that *M. melolontha* exerts positive selection pressure on *T. officinale* defenses (Huber et al. 2016b). Here, we tested the ecological and evolutionary impact of *M. melolontha* root herbivory on seed dispersal and fitness in *T. officinale*. We planted offspring of sympatric and allopatric *T. officinale* populations into the field and selectively infested them with *M. melolontha* larvae. We then assessed the morphological seed characteristics that determine *T. officinale* seed dispersal capacity by linking vertical seed fall rate with seed morphology. We compared dispersal and fitness related traits of seeds collected from plants under grub attack with traits of seeds collected from non-attacked plants. By comparing grub-attack induced changes in seed dispersal and fitness of *T. officinale* populations with different evolutionary histories, we present evidence that *M. melolontha* affects plant fitness and dispersal through induced phenotypic plasticity that is shaped by natural selection.

## Material and Methods

### Morphological seed characteristics underlying T. officinale seed dispersal capacity

Average horizontal dispersal distance (x) of wind-dispersed seeds can be calculated using a common ballistic model by multiplying the shedding height of the seed (*H*) with the horizontal wind speed (*u*), divided by the vertical fall rate of the seed (F): *x* = *Hu*/F (Pasquill & Smith 1983; Brock et al. 2005). Seed fall rate is determined by seed traits such as seed mass and seed shape (Matlak 1987; Meyer & Carlson 2001; Skarpaas et al. 2011). To identify the morphological seed traits that contribute most strongly to seed dispersal potential of *T. officinale*, we measured the fall rate of individual *T. officinale* seeds along with morphological characteristics of individual seeds and used this data to construct a model that predicts dispersal potential from seed morphology. To cover a broad range of genotypes and environments, seeds of a total of 20 plants from the Botanical Garden of the University of Bern, Switzerland (46.95 °N, 7.43 °E, 520 m above sea level), a semi-natural grassland in Boltigen, Switzerland (46.38 °N, 7.23 °E, 1038 m above sea level), and the greenhouse (genotype A34, Verhoeven et al. 2010) were collected. Five seeds per plant were randomly selected for experiments.

To measure seed fall rate, we constructed a semi-automated time-of-flight system (Fig. 1a). The system consists of a plexiglass tube (2 m height, 20 cm diameter, ABC-Kunststoff-Technik GmbH, Germany) that is coated with a fine metal wire grid on the inside to prevent static charges (Fig. 1a). A custom-built sampler on top of the tube ensures standardized, serial release of individual seeds into the time-of-flight tube. The sampler consists of a conveyor belt with 10 seed-containers, an electromotor to move the belt and a series of controllers to the sampler (components from Tinkerforge, Germany). A remote control allows the user to release a seed and simultaneously start a digital timer by pressing the start button. As soon as the seed crosses the finish line at the bottom of the tube, the user presses the stop button, which stops the timer and saves the time-of-flight in a small memory module. The source code of the control software and a list of the individual components can be found on Github (to be inserted at a later time). Using this system, 100 test seeds were measured three times, and average fall rates for each seed were calculated. As physiological parameters that may determine fall rate, seed weight, length and width, achene length, pappus stem length and pappus hair length were determined. To standardize measures, a glass plate was put on top of the seed, and traits were measured two-dimensionally from the top. Linear regression models were then built using falling rate as response variable and combinations of seed traits as explanatory variables. We then compared the models using ANOVAs to select the best-performing model. Major variables contributing to fall rate (seed mass and pappus hair length, see results), were selected as measures of seed dispersal potential in subsequent experiments.

**Fig. 1.**
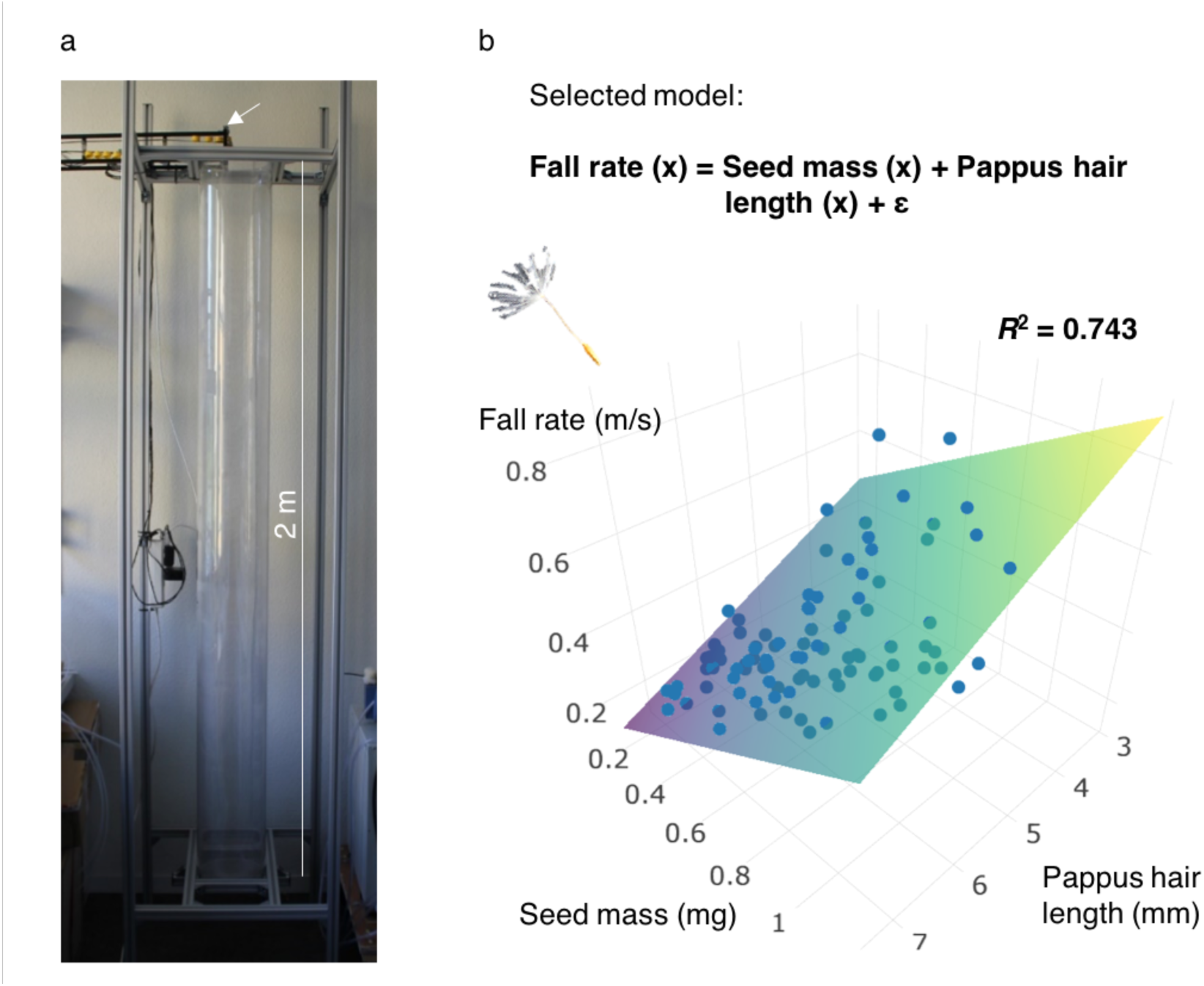
Seed mass and pappus hair length determine dispersal potential of dandelion seeds. (a) Photograph of a custom-built plexiglass time-of-flight tube with semi-automatic seed dispenser (white arrow) to determine fall rates of individual dandelion seeds. (b) A linear model predicts seed fall rate using seed mass and pappus hair length. The adjusted *R*^2^ of the model-fit is displayed.

### Effect of root herbivory on seed dispersal and fitness

To examine how root herbivory affects seed dispersal and fitness of allopatric and sympatric *T. officinale* plants, we performed a one-year experiment in a semi-natural grassland using seeds from 18 *T. officinale* populations evolved in the presence or absence of *M. melolontha* over the last 20 years (Table S2). A detailed description of the evolutionary history of the majority of these populations can be found in Huber et al. (2016b). In brief, nine populations originated from areas in Switzerland and Germany with high documented *M. melolontha* densities over the last 20 years (sympatric populations) and nine populations came from areas with no or only low *M. melolontha* abundance over the last 20 years (allopatric populations). Within each type of evolutionary history, six populations consisted of sexual, outcrossing diploids, and three populations consisted of apomictic, clonal triploids. Triploid seeds were collected directly from their respective mother plants. Diploid seeds were obtained by hand-pollination within populations. For each population, we created a seed mixture by mixing 15 seeds per plant from six randomly selected plants, giving a total of 90 seeds per population. Seed mixtures were germinated in the greenhouse and after two weeks, seedlings were individually transplanted into plastic pots (5 x 5 x 5 cm) filled with soil (Selmaterra, Bigler Samen AG, Thun, Switzerland). Three months after germination, in March 2016, the plants were introduced to the field site in Boltigen (46.38 °N, 7.23 °E, 1038 m above sea level). The field site was established in a semi-natural, organically managed grassland. *T. officinale* occurs naturally at the field site. *M. melolontha* is present in low abundance in this region.

For the experiment, a plot of 50 m x 15 m was fenced in to prevent cattle from entering. Plants were transplanted individually into bigger pots (13 x 13 x 13 cm) filled with soil (Selmaterra, Bigler Samen AG, Thun, Switzerland). To enable exchange of exudates, a hole was cut into the bottom of the pots (5 cm diameter) and a permeable root barrier (15 x 15 cm; Trenn-Vlies, GeoTex Windhager, Switzerland) was put into the pots before transplanting. Then we excavated soil blocks (approximately 14 x 14 x 14 cm) in the field plot in seven parallel rows, with 40 cm space between the holes and 210 cm space between the rows and inserted the plants into the field in a fully randomized design (Fig. S1). This setup allowed us to embed the plants into their natural environment while controlling root herbivore exposure. To infest plants with *M. melolontha*, individual second instar larvae were introduced to half of the pots in a randomized fashion in the beginning of June 2016. *M. melolontha* larvae were collected from a field in Uri, Switzerland (46.45 °N, 8.42 °E, 822 m above sea level) and reared on carrots for 2 weeks prior to the start of infestation.

### Measurement of seed dispersal and fitness

On the northern hemisphere, *T. officinale* has its main flowering peak in spring and a secondary flowering peak in late summer / early autumn (Stewart-Wade et al. 2002). We continuously monitored flower development during both flowering peaks (summer 2016 and spring 2017). Every 3-4 days, all fully ripened seeds were collected for analysis of dispersal-related seed traits. For collection, seeds of inflorescences were carefully transferred into a small paper bag without cutting or damaging the flower and flower stem heights were measured. If seeds of an inflorescence were missing from the flower heads, we estimated the percentage of remaining seeds. Seeds were stored in darkness at 4 °C until analysis.

For seed phenotyping, one seed head from summer 2016 and one seed head from spring 2017 were randomly chosen from each plant. Five to twelve seeds with no visible signs of damage were randomly selected from the different seed heads and phenotyped. Average seed weight and average pappus hair length per seed head were then determined. Seed fall rate was calculated as a function of seed mass and length of pappus hairs. Seed fall rate, vertical wind speed and shedding height of the seed then allowed for an estimation of the dispersal potential for each capsule. A fixed value of 23.88 cm, corresponding to the average stem height in summer 2017, was chosen as a release height for our calculations. We used a constant release height, as the variation in stem height between genotypes recorded in summer 2017 was minimal compared to the variation in seed traits (Table S1, Fig. S2). For wind speed, we used 2 m/s, which is the yearly average wind speed in 2016 recorded by the nearest SwissMetNet surface weather station (Boltigen, 46.37 °N, 7.23 °E, 820 m above sea level).

To determine if the dispersal potential and the number of viable offspring per plant are correlated, we measured germination rates of collected seeds in a climate chamber in August 2017 (temperature at day 22 °C and at night 18 °C; 16 h light and 6 h darkness; 65% humidity; light intensity approximately 120 μmol*m^-2^*s^-1^). From each phenotypic analysed seed capsule, ten seeds were randomly selected and equally distributed in a small pot (5.5 cm diameter) filled with moist seedling substrate. The pots were watered regularly and checked daily. Each day we recorded the number of germinated seeds per pot and after 14 days, germination rate was calculated by dividing the number of emerged seedlings through the number of seeds. By multiplying germination rate with the number of flowers per plant and number of seeds per head per plant, we then estimated the number of viable offspring per plant as a proxy for plant fitness (Erb 2018).

### Measurement of vegetative growth

For each of the 797 plants in the field experiment, we determined plant vegetative growth by counting the number of leaves and measuring the length of longest leaf as a proxy for plant performance (Huber et al. 2016a). Plant performance was determined on a monthly basis from June 2016 to June 2017, with the exception of January-March 2017, when snow cover prevented measurements. In mid-July 2016, the field was mown according to normal agricultural practice. Before mowing, we individually collected the aboveground biomass of each plant. Shoots were cut 5 cm above ground, put into paper bags, dried in an oven at 80 °C until constant mass was reached and weighted. In June 2017, the experiment was terminated, and aboveground biomass was measured again.

### Statistical analyses

All statistical analyses were performed in R 3.3.3 (R Core Team 2017). To predict seed fall rate, ‘Seed mass’ and ‘Pappus hair length’ were used in a linear regression model. No interactions were included in the model. The predictive power of the model was evaluated using k-fold cross validation. For the linear regression model, the function ‘lm’ of the stats package was used (R Core Team 2017) and the validation of the predictive power was done with the function ‘train’ of the ‘caret’ package (Kuhn 2008).

For the analysis of seed traits, mean values per population were calculated separately for plants with and without grub infestation. Mean population values were then examined with linear models. Multiple linear models were built with ‘Herbivory’ x ‘Evolutionary history’ x ‘Ploidy level’ as explanatory variables and each recorded seed parameter as response variable (‘Flower heads’, ‘Seeds per head’, ‘Pappus hair length’, ‘Seed mass’, ‘Dispersal potential’, ‘Germination rate’ and ‘Offspring per plant’). For analysis of vegetative growth, a linear model was employed on ‘Biomass’ as response variable with ‘Herbivory’ x ‘Evolutionary history’ x ‘Ploidy level’ as explanatory variables. Further, ‘Length of longest leaf’ was analysed as response variable in two linear models with ‘Herbivory’ x ‘Evolutionary history’ x ‘Time’ as explanatory variables separately for diploid and triploid plants. All models were examined using the function ‘plotresid’ (package ‘RVAMemoire’ (Hervé 2018)). Significance of explanatory variables was tested using the function ‘Anova’ (package ‘car’ (Fox & Weisberg 2011)) and results were plotted using the package ‘ggplot2’ (Wickham 2009). Similar results were obtained with models for values of individual plants, which included ‘Population’ as random factor, compared to models for mean values per population.

To investigate if changes in seed mass are linked to changes in biomass, correlation strength between ‘Seed mass’ and Biomass’ was analysed separately for seeds from summer 2016 and spring 2017. Further, to examine if there is a trade-off regarding the plant’s investment into seed production optimized for dispersal or germination, correlation strength between ‘Dispersal potential’ and ‘Germination rate’ was analysed separately for seeds from summer 2016 and spring 2017.

To investigate direct and indirect relationships between and within analysed seed and growth traits, we used structural equation modelling (SEM) (Shipley 2016). We developed an a priori model based on the results of our data analysis and on physiological and ecological knowledge, which includes all measured plant traits except the estimated variables ‘Dispersal potential’ and ‘Offspring per plant’, as those were calculated out of other seed traits. We hypothesized direct effects of season (summer 2016 vs. spring 2017) on the measured seed and flower traits and on biomass production. Further, we assumed direct effects of biomass on the measured seed and flower traits and an indirect effect of biomass on germination rate through seed mass and pappus hair length. In addition, we hypothesized an indirect effect of number of flower heads per plant and number of seeds per head on germination rate through seed mass and pappus hair length. We assumed bidirectional links between seed mass and pappus hair length and between number of flower heads per plant and number of seeds per head. To avoid large differences in the variances among the variables we first scaled all variables to an equal range by dividing through powers of 10 if necessary. We then used the R package ‘lavaan’ (Rosseel 2012) for fitting and testing the SEM. Model fit indices (CFI = 0.999, TLI = 0.996, RMSEA = 0.018) indicated good model fit, so maximum likelihood was used to estimate standardized path coefficients.

## Results

### Seed weight and pappus hair length determine dispersal potential of T. officinale seeds

Comparisons of multiple linear regression models revealed that vertical seed fall rate of *T. officinale* – and subsequently seed dispersal potential - is predicted well by seed mass combined with pappus hair length. The final model (Fall rate (x) = Seed mass (x) + Pappus hair length (x) + ε) showed high predictive power using these two parameters (R^2^ = 0.743, Fig. 1b). While seed mass increases fall rate and thus decreases dispersal potential, the length of the pappus hair decreases fall rate and thus increases dispersal potential.

### Root herbivory attack increases seed dispersal potential in sympatric populations

During the summer flowering period, *M. melolontha* infestation did not affect pappus hair length (Fig. 2a), but reduced the mass of *T. officinale* seeds (Fig. 2b), resulting in increased dispersal potential compared to control plants (Fig 2c). This response was observed in sympatric, but not allopatric populations (LM, factor ‘EH x H’, *P* = 0.001, Fig. 2b; respectively LM, factor ‘EH x H’, *P* = 0.037, Fig. 2c). During the spring flowering period, no effect of root herbivory on seed traits was detected (Table S1).

**Fig. 2.**
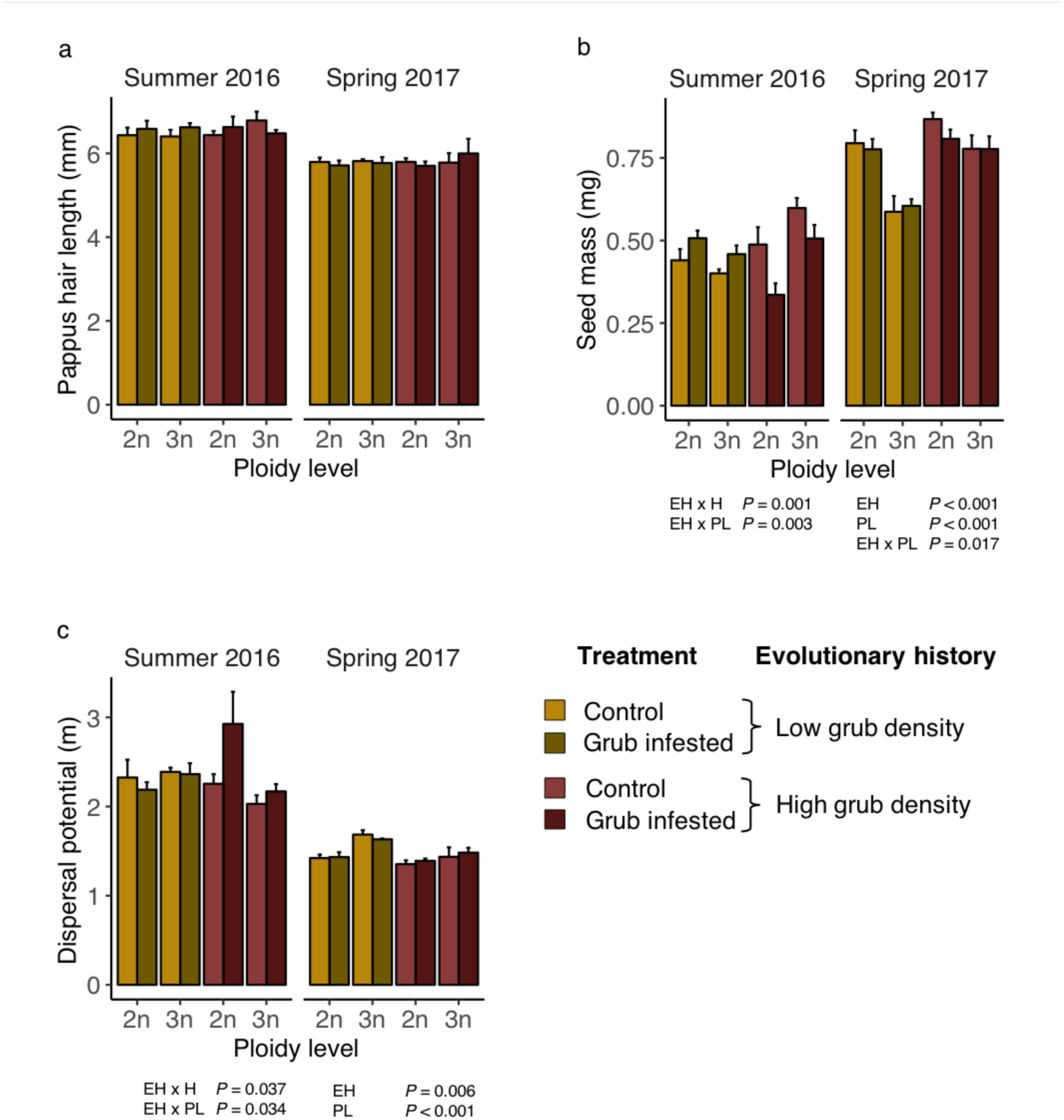
Root herbivore attack increases seed dispersal potential in sympatric dandelion populations. Compared to allopatric populations (yellow bars), sympatric populations (red bars) developed lighter seeds with increased dispersal potential under root herbivory. Seed mass (a), pappus hair length (b) and dispersal potential (c) of seeds collected from plants with (hatched bars) and without (plain bars) root herbivory by *M. melolontha* are shown. Mean values of population means (N = 3-6 populations) and standard errors (± SE) are indicated. *P*-values from linear models are shown (significant effects only). EH: Evolutionary history of population. H: Root herbivory treatment. PL: Ploidy level. 2n: Diploid populations. 3n: Triploid populations.

### Evolutionary history and cytotype interact to determine seed dispersal potential

We detected significant effects of evolutionary history and cytotype on seed mass and dispersal potential (Fig. 2). During the summer flowering period, seeds of sympatric triploids were heavier and had a lower dispersal potential than seeds of sympatric diploids, while the opposite was the case for allopatric populations (LM, factor ‘EH x PL’, *P* = 0.003, Fig. 2b; respectively LM, factor ‘EH x PL’, *P* = 0.034, Fig. 2c). During the spring flowering period, seeds of sympatric populations were heavier and had a lower dispersal potential than seeds of allopatric populations (LM, factor ‘EH’, *P* < 0.001, Fig. 2b; respectively LM, factor ‘EH’, *P* = 0.006, Fig. 2c), and triploid seeds were generally lighter and thus had a higher dispersal potential than diploid seeds (LM, factor ‘PL’, *P* < 0.001, Fig. 2b; respectively LM, factor ‘PL’, *P* < 0.001, Fig. 2c).

### Root herbivory reduces plant fitness

During the summer flowering period, root herbivory did not significantly affect the number of flower heads, the number of seeds or the percentage of germinating seeds per plant (Fig. 3a-c). The total number of offspring per plant, which was calculated by multiplying flower heads, seeds per head and germination rate, however, was reduced by *M. melolontha* attack in sympatric, but not in allopatric populations (LM, factor ‘EH x H’, *P* = 0.039, Fig. 3d). In spring, *M. melolontha* reduced the number of flower heads and total viable offspring per plant independently of the other factors (LM, factor ‘H’, *P* < 0.001, Fig. 3a; respectively LM, factor ‘H’, *P* = 0.003, Fig. 3d). Evolutionary history and cytotype also affected plant fitness traits (Fig. 3). Triploid seeds from the summer flowering period germinated better, leading to a higher number of offspring per plant (LM, factor ‘PL’, *P* =0.002, Fig. 3c; respectively LM, factor ‘PL’, *P* < 0.001, Fig. 3d). During the spring flowering period, triploids produced more flower heads per plant (LM, factor ‘PL’, *P* = 0.001, Fig. 3a), but fewer seeds per flower head (LM, factor ‘PL’, *P* = 0.002, Fig. 3b), and showed no difference in germination rate and offspring per plant (Fig. 3c-d). Sympatric populations produced fewer flowers than allopatric populations during the spring flowering period, and the number of viable offspring was lower for sympatric populations than allopatric populations (LM, factor ‘EH’, *P* < 0.001, Fig. 3a; respectively LM, factor ‘EH’, *P* = 0.017, Fig. 3d).

**Fig. 3.**
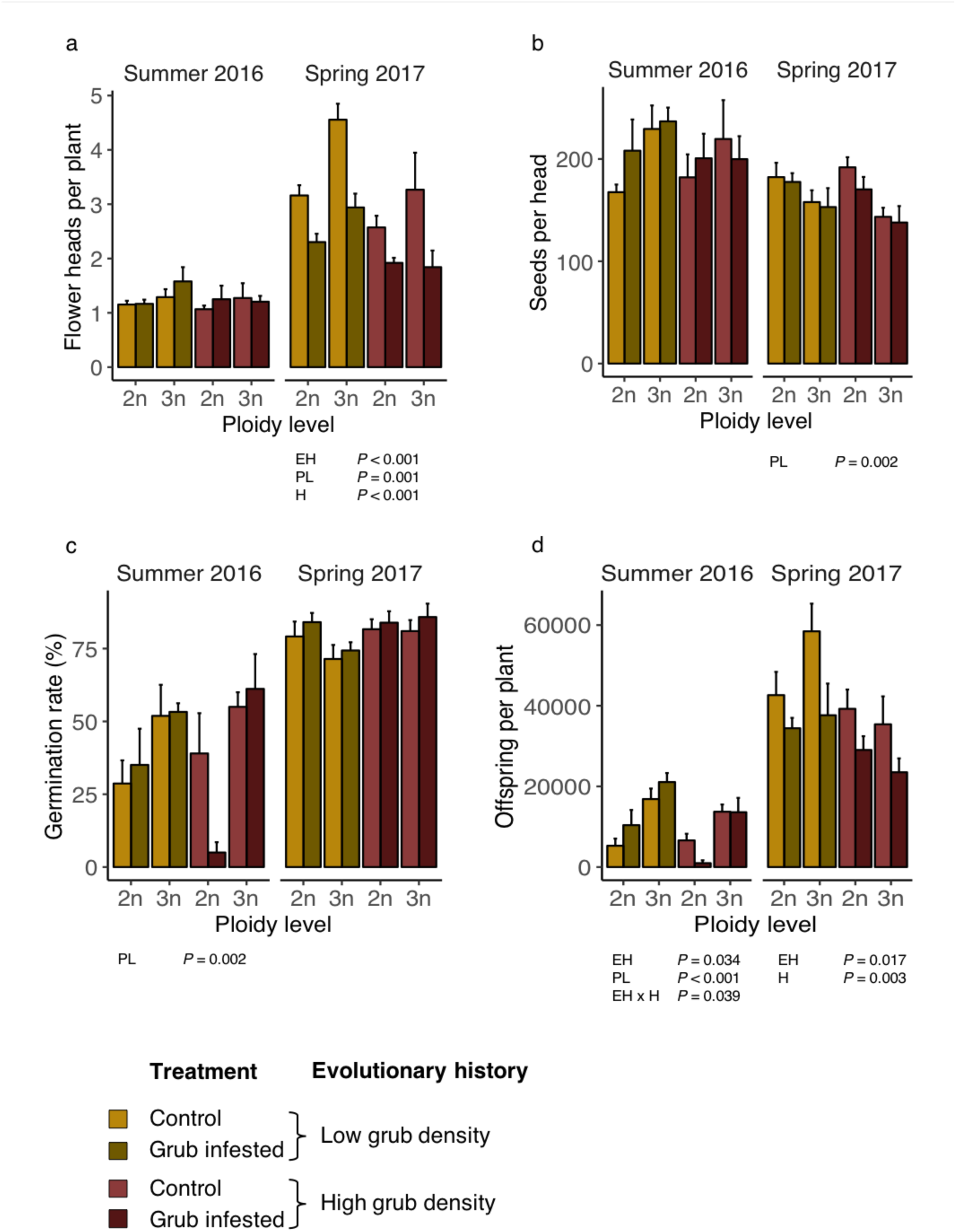
Root herbivore attack reduces plant fitness. Flower heads per plant (a), seeds per head (b), germination rate (c) and offspring per plant (d) of plants with (hatched bars) and without (plain bars) root herbivory by *M. melolontha* are shown. Mean values of population means (N = 3-6 populations) and standard errors (± SE) are indicated. *P-*values from linear models are displayed (significant effects only). EH: Evolutionary history of population. H: Root herbivory treatment. PL: Ploidy level. 2n: Diploid populations. 3n: Triploid populations.

### Root herbivory decreases vegetative growth

To determine whether reduced plant performance upon *M. melolontha* attack may explain the changes in seed mass and subsequently in dispersal potential in sympatric populations, we measured the impact of *M. melolontha* attack on shoot growth and biomass and then correlated shoot biomass with seed mass. The length of the longest leaf, a proxy for biomass accumulation (Huber et al. 2016a) varied through the season and was significantly reduced by *M. melolontha* infestation (Fig. 4a). Shoot biomass was significantly reduced in plants attacked by *M. melolontha* during both sampling periods (LM, factor ‘Herbivory’, *P* = 0.009 and LM, factor ‘Herbivory’, *P* = 0.047, Fig. 4b). Further, during the summer sampling period, triploids produced more biomass than diploids (LM, factor ‘Ploidy level’, *P* < 0.001, Fig. 4b). However, none of these effects was different between sympatric and allopatric populations, suggesting that the increase in seed dispersal capacity upon *M. melolontha* attack in sympatric populations is not due to a general plant stress response.

**Fig. 4.**
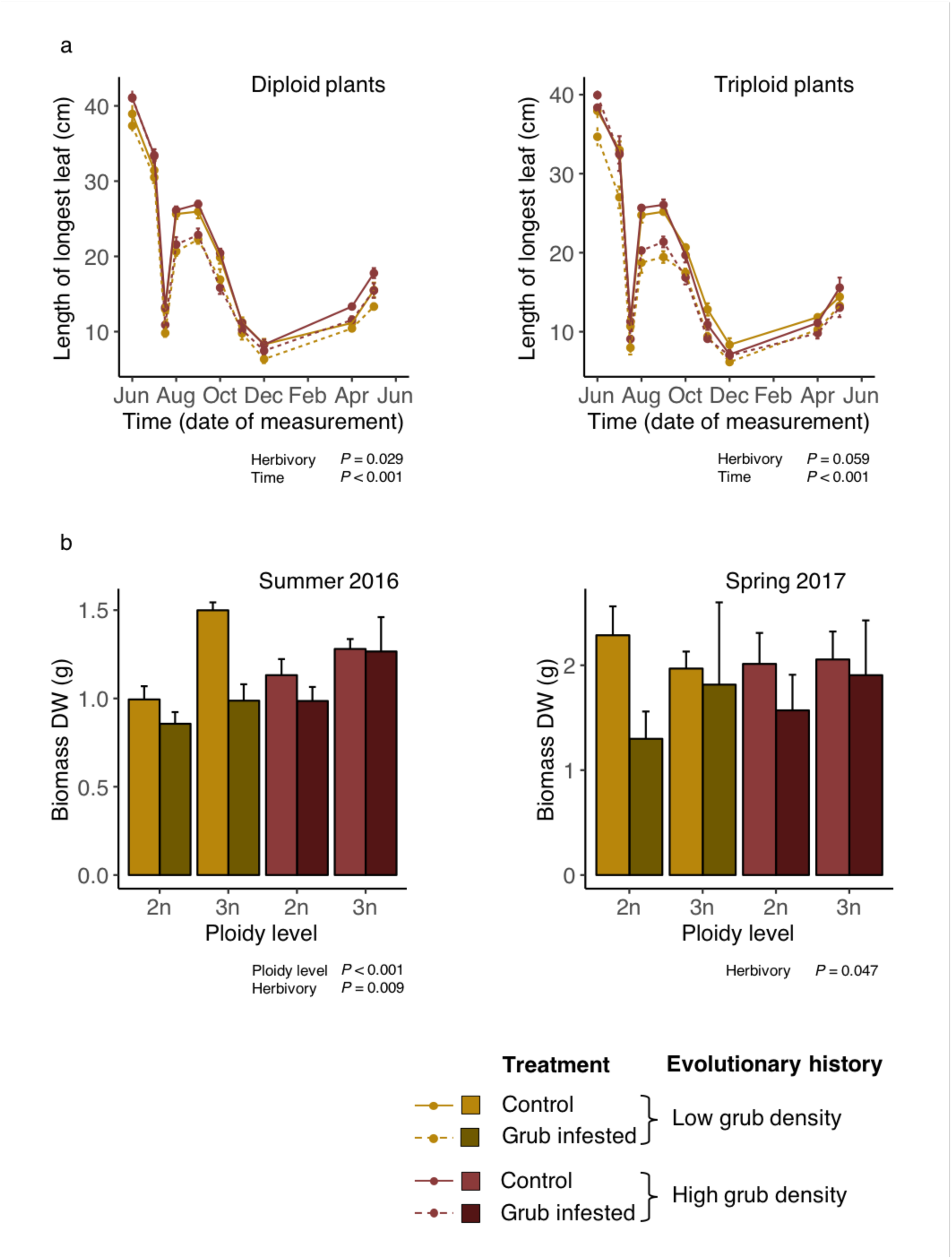
Rood herbivore attack reduces vegetative growth. (a) Length of longest leaf in diploid and in triploid populations under infestation with *M. melolontha* (dashed lines) compared to plants without root herbivory (full lines). Vegetation was cut back at the end of July. (b) Above ground biomass decreased under root herbivory (hatched bars) compared to biomass of plants without root herbivory (plain bars). Mean values of population means (N = 3-6 populations) and standard errors (± SE) are shown. *P-*values from linear models are displayed (significant effects only). 2n: Diploid populations. 3n: Triploid populations.

### Correlations between vegetative growth, fitness and dispersal

Structural equation modelling (SEM) revealed that the sampling period (‘Season’) had an influence on both vegetative and generative traits. In summer 2016, *T. officinale* plants produced more seeds per head and seeds with longer pappus hair. In spring 2017, plants produced more biomass, more flower heads per plant, heavier seeds with higher germination rate, but shorter pappus hair (Fig. 5a).

**Fig. 5.**
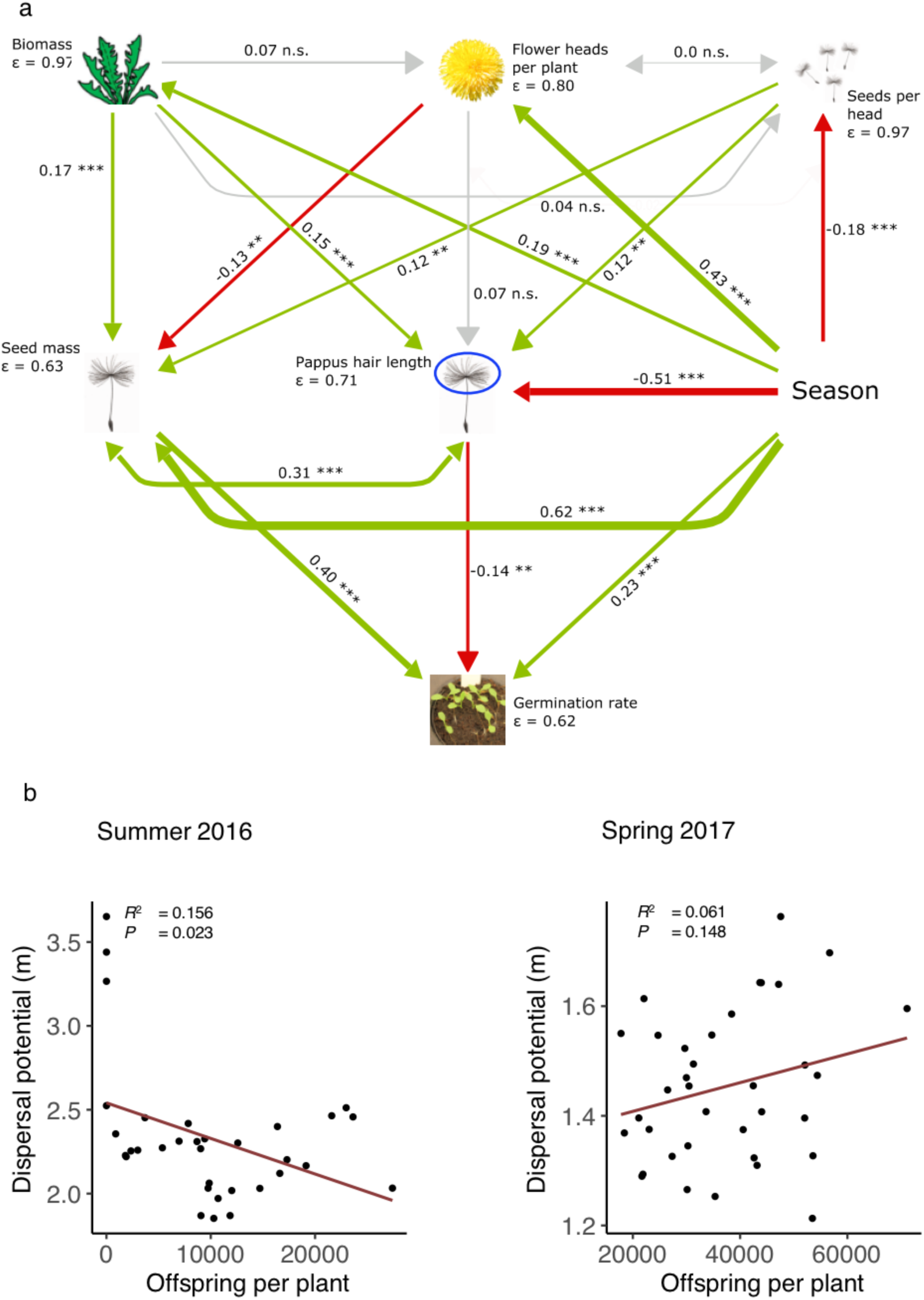
Dispersal and establishment are negatively correlated during the summer flowering period. (a) Structural equation model showing pathways between phenotypic plant parameters. Positive links are indicated by green arrows and negative links are indicated by red arrows. Grey arrows represent non-significant links. Red arrows coming from ‘Season’ represent stronger effects of sampling period ‘summer 2016’, whereas green arrows coming from ‘Season’ represent stronger effects of sampling period ‘spring 2017’. Line thicknesses of arrows correspond to the effect size given by their standardized path coefficients, which are shown next to each path. Asterisks next to path coefficients indicate *P*-values (\*\*\**P* < 0.001; \*\**P* < 0.01; \**P* < 0.05; n.s. *P* ≥ 0.05). Residual variance (ε) is given for all response variables. (b) Strength of correlation between dispersal potential and offspring per plant for seeds collected in summer 2016 and in spring 2017. Linear regression lines of population means and corresponding *R*^2^- and *P-*values are shown.

Across all plants, vegetative biomass production explained seed mass and pappus hair length (Fig. 5a). However, across populations and analysed for each sampling season separately, shoot biomass was not correlated with average seed mass and hence dispersal potential (*R*^2^ = 0.009, *P* = 0.593 and *R*^2^ = 0.080, *P* = 0.095, Fig. S3). Seed mass explained pappus hair length and germination rate. (Fig. 5a). Pappus hair length was a negative predictor of germination rate. The number of flowers per head was a negative predictor of seed mass, while the number of seeds per head was a positive predictor of seed mass and pappus hair length (Fig. 5a).

To further understand the relationship between dispersal potential (calculated from seed mass and pappus hair length) and total offspring per plant (calculated from flower heads, seeds per head and germination rate), we correlated these two parameters. Dispersal potential and offspring per plant were negatively correlated for the summer flowering period, but not the spring flowering period (*R*^2^ = 0.156, *P* = 0.023 and *R*^2^ = 0.061, *P* = 0.148, Fig. 5b). Together, these results show that *T. officinale* dispersal and reproductive potential differ markedly between the summer and spring flowering periods, and that there seems to be a tradeoff between dispersal and reproduction during the summer flowering period, which is mostly driven by seed mass, which increases germination, but decreases dispersal potential.

## Discussion

Plants can escape from unfavourable environments through seed dispersal. How seed dispersal capacity is affected by herbivory, however, is not well understood. Here, we show that *T. officinale* plants from sympatric, but not allopatric populations, increase their seed dispersal potential upon root herbivore attack in the field by reducing seed mass at the expense of the number of viable offspring. Below, we discuss the mechanisms and ecological implications of these findings.

Whether seed dispersal is an effective strategy to escape from hostile environments depends on the distance that can be covered by the dispersing seeds relative to the size of the unfavorable habitat. Attack by root-feeding *M. melolontha* larvae resulted in increased seed dispersal potential in sympatric *T. officinale* plants during the summer flowering period. Assuming an average wind speed of 2m/s, dispersal potential for grub-attacked sympatric populations can be estimated to exceed dispersal of non-attacked populations by 41 cm. As third instar larvae, *M. melolontha* can move up to 20 cm per day (Hasler 1986). An increased seed dispersal distance of 41 cm may therefore not be sufficient to escape a patch with high *M. melolontha* density. However, chances for long-distance dispersal induced by vertical wind turbulence are also higher for seeds with lower falling velocity. Although long-distance dispersal is relatively rare, producing seeds with lower fall rates does enhance the probability of *T. officinale* offspring to be dispersed over hundreds of meters under favorable conditions, which may allow the plant’s offspring to escape from heavily root herbivore infested patches.

Theory predicts that increased seed dispersal is advantageous under locally stressful conditions and may therefore be adaptive (Levin et al. 2003, Ronce 2007). To which extend the herbivory-induced changes in seed traits observed here represents a form of adaptive plasticity is not fully clear. Adaptive plasticity, defined as plasticity maintained through natural selection, should lead to increased fitness (Miner et al. 2005). However, dispersal distance *per se* is not a direct fitness measurement, as the benefit of dispersal depends on the environmental context (van Kleunen & Fischer 2005; Teller et al. 2014). In this context, it is worth noting that we observed an increase in seed dispersal potential during grub attack in sympatric *T. officinale* populations, which have evolved in the presence of *M. melolontha* over the last decades, but not in allopatric populations that have not been under white grub pressure. This result supports the notion that seed dispersal traits are subject to natural selection (Cheptou et al. 2008; Ronce 2007; García-Verdugo et al. 2017; Saastamoinen et al. 2018) and indicates that root herbivores may exert selection pressure on seed phenotypic plasticity.

In general, the seed dispersal capacity of a plant may increase under unfavorable environmental conditions due to “stress flowering”, leading to suboptimal seed development and, thus, lighter seeds (Germain & Gilbert 2014, Begcy & Walia 2015). Our results suggest, however, that physiological stress is unlikely to explain the increase in dispersal potential in root-herbivore attacked dandelions. We found that *M. melolontha* attack reduces growth and biomass accumulation of dandelion plants, but that these effects are similar in sympatric and allopatric populations. Thus, we expect that the extent of stress that is imposed by root herbivory does not differ between allopatric and sympatric populations. As only sympatric populations adjust their seed phenotype upon root herbivore attack, we, therefore, propose that this change represents a form of herbivore-induced plant response rather than a secondary effect of physiological stress.

Seed mass influences dispersal potential and germination ability in opposite directions and therefore represents a physiological trade-off. Many studies empirically support this assumption (e.g. Soons & Heil 2002; Jakobsson & Eriksson 2003). Our study confirms that vertical falling velocity and hence distribution distance of *T. officinale* seeds as well as germination rate are strongly linked to seed mass. The capacity to disperse seeds thus comes with a reduction in total viable offspring. Thus, dandelions may trade establishment ability against higher dispersal potential under root herbivore attack. The high cost of higher dispersal likely contributed to the inducibility of this phenomenon. Interestingly we found a significant negative relationship between dispersal distance and number of viable offspring for the first sampling period in summer, when root herbivory had a significant effect on dispersal, but no significant link in the second sampling period, where root herbivory did not change seed phenotypes. Thus, the negative relationship between dispersal and seed mass seems to be context dependent and more strongly expressed under environmental stress.

Herbivores exert selection pressure on plants, leading to the evolution of tolerance and defense strategies (Núñez-Farfán et al. 2007). Earlier work demonstrated that sympatric dandelion populations that grow in the presence of *M. melolontha* produce higher amounts of repellent root sesquiterpene lactones, which may benefit them by becoming less attractive to *M. melolontha* (Huber et al. 2016b). The experimental setup in this study controlled for these potential effects by confining root herbivores to individual dandelion plants. Our results suggest that, when *M. melolontha* larvae do not have a choice between different plants, there is no direct benefit to producing the repellent chemicals. On the contrary, sympatric populations suffered from the same biomass reduction and did not produce more flowers or seeds under root herbivore attack than allopatric populations. In combination with our earlier studies (Huber et al. 2016a, Huber et al. 2016b, Bont et al. 2017), these results suggest that high herbivore pressure selects for repellent root chemicals and higher dispersal potential, but not stronger resistance or tolerance in dandelion. These results emphasize the need to evaluate the evolution of plant defense strategies in a community context (Agrawal et al. 2012, Poelman & Kessler 2016) and to include seed and dispersal traits into these analyses (Erb 2018).

Through feeding on roots, belowground insects can severely impair plant performance. From an evolutionary point of view, presence of root-feeders may therefore act as selective force on plant traits involved in seed distribution and establishment. In our study, exposure to *M. melolontha* resulted in reduced flower production, but it also resulted in the production of better dispersible seeds - depending on the sampling period and the evolutionary history of *T. officinale.* Thus, our findings suggest an important, but context-dependent influence of root herbivory on plant reproduction and dispersal traits.

## Acknowledgements

We thank Miguel Salinas, Robin Bautzmann, Conradin Lutz and Gabriel Zala for their help with seed phenotyping. We are grateful to the members of the Biotic Interactions Group at the Institute of Plant Sciences (University of Bern) as well as to Zephyr Züst, Cyrill Delfgou, Cédric Zahnd, Célia Ruiz, Armin Komposch, Valentin Pulver, Tala Bürki and Andrea Bonini for their help during field work. This study was supported by the Swiss National Science Foundation (Grant No. 153517) and the Seventh Framework Programme for Research and Technological Development of the European Union (FP7 MC-CIG 629134).

## Supporting information

Additional supporting information may be downloaded via the online version of this manuscript.

## References

Agrawal, A.A., Hastings, A.P., Johnson, M.T.J., Maron, J.L. & Salminen, J.-P. (2012). Insect herbivores drive real-time ecological and evolutionary change in plant populations. Science, 338, 113–116.

Begcy, K. & Walia, H. (2015). Drought stress delays endosperm development and misregulates genes associated with cytoskeleton organization and grain quality proteins in developing wheat seeds. Plant Sci., 240, 109–119.

Bont, Z., Arce, C., Huber, M., Huang, W., Mestrot, A., Sturrock, C.J., et al. (2017). A herbivore tag-and-trace system reveals contact-and density-dependent repellence of a root toxin. J. Chem. Ecol., 43, 295–306.

Brock, M.T., Weinig, C. & Galen, C. (2005). A comparison of phenotypic plasticity in the native dandelion Taraxacum ceratophorum and its invasive congener T. officinale. New Phytol., 166, 173–183.

Cheptou, P.O., Carrue, O., Rouifed, S. & Cantarel, A. (2008). Rapid evolution of seed dispersal in an urban environment in the weed Crepis sancta. Proc. Natl. Acad. Sci., 105, 3796–3799.

Cummins, C., Seale, M., Macente, A., Certini, D., Mastropaolo, E., Viola, I.M., et al. (2018). A separated vortex ring underlies the flight of the dandelion. Nature, 562, 414–418.

de la Peña, E. & Bonte, D. (2014). Above-and belowground herbivory jointly impact defense and seed dispersal traits in Taraxacum officinale. Ecol. Evol., 4, 3309–3319.

de la Peña, E., D’hondt, B. & Bonte, D. (2011). Landscape structure, dispersal and the evolution of antagonistic plant-herbivore interactions. Ecography, 34, 480–487.

Erb, M. (2018). Plant defenses against herbivory: Closing the fitness gap. Trends Plant Sci., 23, 187–194.

Fox, J. & Weisberg, S. (2011). An {R} companion to applied regression. Second edition. Sage, Thousand Oaks CA.

García-Verdugo, C., Mairal, M., Monroy, P., Sajeva, M. & Caujapé-Castells, J. (2017). The loss of dispersal on islands hypothesis revisited: Implementing phylogeography to investigate evolution of dispersal traits in Periploca (Apocynaceae). J. Biogeogr., 44, 2595–2606.

Germain, R.M. & Gilbert, B. (2014). Hidden responses to environmental variation: Maternal effects reveal species niche dimensions. Ecol. Lett., 17, 662–669.

Greene, D.F. & Quesada, M. (2005). Seed size, dispersal, and aerodynamic constraints within the Bombacaceae. Am. J. Bot., 92, 998–1005.

Hasler, T. (1986). Abundanzund Dispersionsdynamik von Melolontha Melolontha (L.) in Intensivobstanlagen. Eidgenössische Technische Hochschule Zürich.

Hauss, R. (1975). Methoden und erste Ergebnisse zur Bestimmung der Wirtspflanzen des Maikäferengerlings (Melolontha melolontha L.). Mitteilungen aus der Biol. Bundesanstalt für Landund Forstwirtschaft Berlin Dahlen, 73–77.

Hauss, R. & Schütte, F. (1976). Zur Polyphagie der Engerlinge von Melolontha melolontha L. an Pflanzen aus Wiese und Ödland. Anzeiger für Schädlingskd., 49, 129–132.

Hervé, M. (2018). Package “RVAideMemoire.”

Huber, M., Bont, Z., Fricke, J., Brillatz, T., Aziz, Z., Gershenzon, J., et al. (2016b). A belowground herbivore shapes root defensive chemistry in natural plant populations. Proc. Biol. Sci., 283.

Huber, M., Epping, J., Schulze Gronover, C., Fricke, J., Aziz, Z., Brillatz, T., et al. (2016a). A latex metabolite benefits plant fitness under root herbivore attack. PLoS Biol., 14, 1–27.

Jakobsson, A. & Eriksson, O. (2003). Trade-offs between dispersal and competitive ability: A comparative study of wind-dispersed Asteraceae forbs. Evol. Ecol., 17, 233–246.

Kawecki, T.J. & Ebert, D. (2004). Conceptual issues in local adaptation. Ecol. Lett., 7, 1225–1241.

van Kleunen, M. & Fischer, M. (2005). Constraints on the evolution of adaptive phenotypic plasticity in plants. New Phytol., 166, 49–60.

Kuhn, M. (2008). Building predictive models in R using the caret package. J. Stat. Softw., 28, 1–26.

Lau, J.A., McCall, A.C., Davies, K.F., McKay, J.K. & Wright, J.W. (2008). Herbivores and edaphic factors constrain the realized niche of a native plant. Ecology, 89, 754–762.

Leishman, M.R. & Westoby, M. (1994). Hypotheses on seed size: Tests using the semiarid flora of Western New South Wales, Australia. Am. Nat., 143, 890–906.

Levin, S.A., Muller-Landau, H.C., Nathan, R. & Chave, J. (2003). The ecology and evolution of seed dispersal: A theoretical perspective. Annu. Rev. Ecol. Evol. Syst., 34, 575–604.

Martorell, C. & Martínez-López, M. (2014). Informed dispersal in plants: Heterosperma pinnatum (Asteraceae) adjusts its dispersal mode to escape from competition and water stress. Oikos, 123, 225–231.

Matlack, G.R. (1987). Diaspore size, shape, and fall behavior in wind-dispersed plant species. Am. J. Bot., 74, 1150–1160.

Meyer, S.E. & Carlson, S.L. (2001). Achene mass variation in Ericameria nauseosus (Asteraceae) in relation to dispersal ability and seedling fitness. Funct. Ecol., 15, 274–281.

Miner, B.G., Sultan, S.E., Morgan, S.G., Padilla, D.K. & Relyea, R.A. (2005). Ecological consequences of phenotypic plasticity. Trends Ecol. Evol., 20, 685–692.

Nathan, R. & Muller-Landau, H.C. (2000). Spatial patterns of seed dispersal, their determinants and consequences for recruitment. Trends Ecol. Evol., 15, 278–285.

Núñez-Farfán, J., Fornoni, J. & Valverde, P.L. (2007). The evolution of resistance and tolerance to herbivores. Annu. Rev. Ecol. Evol. Syst., 38, 541–566.

Pasquill, F. & Smith, F.B. (1983). Atmospheric diffusion. 3rd edn. Ellis Horwood, Chichester.

Poelman, E.H. & Kessler, A. (2016). Keystone herbivores and the evolution of plant defenses. Trends Plant Sci., 21, 477–485.

Poschlod, P., Abedi, M., Bartelheimer, M., Drobnik, J., Rosbakh, S. & Saatkamp, A. (2013). 6.Seed ecology and assembly rules in plant communities. Veg. Ecol. Second Ed., 164–202.

van der Putten, W.H. (2003). Plant defence belowground and spatiotemporal processes. Ecology, 84, 2269–2280.

Van Der Putten, W.H., Vet, L.E.M., Harvey, J.A. & Wäckers, F.L. (2001). Linking above-and belowground multitrophic interactions of plants, herbivores, pathogens, and their antagonists. Trends Ecol. Evol., 16, 547–554.

R Core Team (2017). R: A language and environment for statistical computing. R Found. Stat. Comput. Vienna, Austria.

Ronce, O. (2007). How does it feel to be like a rolling stone? Ten questions about dispersal evolution. Annu. Rev. Ecol. Evol. Syst., 38, 231–253.

Rosseel, Y. (2012). lavaan: An R package for structural equation modelling. J. Stat. Softw., 48, 1–36.

Rutter, M.T. & Rausher, M.D. (2004). Natural selection on extrafloral nectar production in Chamaecrista Fasciculata: The costs and benefits of a mutualism trait. Evolution, 58, 2657–2668.

Saastamoinen, M., Bocedi, G., Cote, J., Legrand, D., Guillaume, F., Wheat, C.W., et al. (2018). Genetics of dispersal. Biol. Rev., 93, 574–599.

Shipley, B. (2016). Cause and correlation in biology: A user’s guide to path analysis, structural equations and causal inference with R. 2nd editio. Cambridge Univ. Press, Cambridge.

Skarpaas, O., Silverman, E.J., Jongejans, E. & Shea, K. (2011). Are the best dispersers the best colonizers? Seed mass, dispersal and establishment in Carduus thistles. Evol. Ecol., 25, 155–169.

Soons, M.B. & Heil, G.W. (2002). Reduced colonization capacity in fragmented populations of wind-dispersed grassland forbs. J. Ecol., 90, 1033–1043.

Stewart-Wade, S.M., Neumann, S., Collins, L.L. & Boland, G.J. (2002). The biology of Canadian weeds. 117. Taraxacum officinale

G. H. Weber ex Wiggers. Can. J. Plant Sci., 825–853.

Tackenberg, O., Poschlod, P. & Kahmen, S. (2003). Dandelion seed dispersal: The horizontal wind speed does not matter for long-distance dispersal - it is updraft! Plant Biol., 5, 451–454.

Teller, B.J., Campbell, C. & Shea, K. (2014). Dispersal under duress: Can stress enhance the performance of a passively dispersed species? Ecology, 95, 2699–2706.

Turnbull, L.A., Coomes, D., Hector, A., Rees, M. (2004). Seed mass and the competition / colonization trade-off?: Competitive interactions and spatial patterns in a guild of annual plants. J. Ecol., 92, 97–109.

Verhoeven, K.J.F., Van Dijk, P.J. & Biere, A. (2010). Changes in genomic methylation patterns during the formation of triploid asexual dandelion lineages. Mol. Ecol., 19, 315–324.

Walter, G.M., Wilkinson, M.J., Aguirre, J.D., Blows, M.W. & Ortiz-Barrientos, D. (2018). Environmentally induced development costs underlie fitness tradeoffs. Ecology, 99, 1391–1401.

Wickham, H. (2009). ggplot2: Elegant graphics for data analysis. Springer-Verlag New York.

Züst, T., Heichinger, C., Grossniklaus, U., Harrington, R., Kliebenstein, D.J. & Turnbull, L.A. (2012). Natural enemies drive geographic variation in plant defenses. Science, 338, 116–120.

