## Supplementary information for "Adapted dandelions increase seed dispersal potential when they are attacked by root herbivores"

### Supporting information

**Table S1** Linear models for seed and growth traits as response variables and root herbivory (H), evolutionary history (EH) and ploidy level (PL) as explanatory variables. *F*- and *P*- values are displayed, with *P*-values < 0.05 in bold.

**Table S2** Description of field sites of *T. officinale* populations used in this study and correspondence to field sites in Huber et al. 2016b.

**Fig. S1** Pictures of the field site in Adlemsried. (a) Field site after the establishment of the plot (March 2016). (b) Field site during the white grub infestation of half of the plants (June 2016).

**Fig. S2** Flower stem height of plants measured in spring 2017 is not influenced by treatment, evolutionary history or ploidy level. Mean values of population means (N = 3-6 populations) and standard errors (± SE) are shown. 2n: Diploid populations. 3n: Triploid populations.

**Fig. S3** Strength of correlation between seed mass and biomass for seeds collected in summer 2016 and in spring 2017. Linear regression lines of population means and corresponding $R^2$- and *P*-values are shown.

#### Table S1

| Response Variable | Factor | Summer 2016 | | Spring 2017 | |
|---|---|---|---|---|---|
| | | *F* | *P* | *F* | *P* |
| Seed mass | Herbivory (H) | 0.77 | 0.39 | 0.94 | 0.34 |
| | Evolutionary history (EH) | 0.16 | 0.69 | 15.88 | **< 0.001** |
| | Ploidy level (PL) | 2.46 | 0.13 | 24.24 | **< 0.001** |
| | H x EH | 13.54 | **0.001** | 0.49 | 0.49 |
| | H x PL | 0.19 | 0.67 | 0.88 | 0.36 |
| | EH x PL | 11.33 | **0.002** | 6.48 | **0.02** |
| | H x EH x PL | 0.40 | 0.53 | 0.05 | 0.82 |
| Pappus hair length | H | 0.47 | 0.50 | 0.10 | 0.76 |
| | EH | 0.19 | 0.67 | 0.09 | 0.77 |
| | PL | 0.16 | 0.70 | 0.73 | 0.40 |
| | H x EH | 0.41 | 0.53 | 0.17 | 0.69 |
| | H x PL | 0.53 | 0.47 | 0.70 | 0.41 |
| | EH x PL | 0.14 | 0.71 | 0.23 | 0.64 |
| | H x EH x PL | 1.07 | 0.31 | 0.44 | 0.51 |
| Dispersal potential | H | 1.41 | 0.25 | 0.18 | 0.68 |
| | EH | 0.51 | 0.48 | 8.76 | **0.006** |
| | PL | 1.50 | 0.23 | 18.50 | **< 0.001** |
| | H x EH | 4.85 | **0.04** | 0.53 | 0.47 |
| | H x PL | 0.51 | 0.48 | 0.13 | 0.72 |
| | EH x PL | 5.05 | **0.03** | 3.81 | 0.06 |
| | H x EH x PL | 1.43 | 0.24 | 0.25 | 0.62 |
| Flower heads | H | 0.83 | 0.37 | 33.57 | **< 0.001** |
| | EH | 0.53 | 0.47 | 17.12 | **< 0.001** |
| | PL | 2.94 | 0.10 | 12.80 | **0.001** |
| | H x EH | 0.01 | 0.91 | 0.33 | 0.57 |
| | H x PL | 0.02 | 0.90 | 4.27 | **0.05** |
| | EH x PL | 0.76 | 0.39 | 3.66 | 0.07 |
| | H x EH x PL | 1.49 | 0.23 | 0.00 | 0.98 |
| Seeds per head | H | 1.02 | 0.32 | 1.36 | 0.25 |
| | EH | 0.12 | 0.74 | 0.21 | 0.65 |
| | PL | 3.29 | 0.08 | 11.61 | **0.002** |
| | H x EH | 0.46 | 0.50 | 0.41 | 0.53 |
| | H x PL | 0.96 | 0.34 | 0.17 | 0.68 |
| | EH x PL | 0.55 | 0.47 | 0.70 | 0.41 |
| | H x EH x PL | 0.01 | 0.94 | 0.18 | 0.67 |
| Germination rate | H | 0.51 | 0.48 | 1.48 | 0.23 |
| | EH | 0.19 | 0.67 | 2.01 | 0.17 |
| | PL | 11.77 | **0.002** | 1.59 | 0.22 |
| | H x EH | 2.32 | 0.14 | 0.04 | 0.85 |
| | H x PL | 1.03 | 0.32 | 0.00 | 0.96 |
| | EH x PL | 0.85 | 0.36 | 2.14 | 0.15 |
| | H x EH x PL | 1.97 | 0.17 | 0.13 | 0.72 |
| Offspring per plant | H | 0.33 | 0.57 | 10.44 | **0.003** |
| | EH | 5.03 | **0.03** | 6.48 | **0.02** |
| | PL | 27.92 | **< 0.001** | 0.40 | 0.53 |
| | H x EH | 4.77 | **0.04** | 0.05 | 0.82 |
| | H x PL | 0.29 | 0.59 | 0.88 | 0.36 |
| | EH x PL | 0.11 | 0.74 | 3.49 | 0.07 |
| | H x EH x PL | 0.65 | 0.43 | 0.51 | 0.48 |
| Biomass | H | 8.01 | **0.01** | 4.03 | **0.05** |
| | EH | 2.35 | 0.14 | 0.01 | 0.91 |
| | PL | 15.16 | **< 0.001** | 0.29 | 0.60 |
| | H x EH | 1.55 | 0.22 | 0.51 | 0.48 |
| | H x PL | 0.78 | 0.38 | 1.09 | 0.30 |
| | EH x PL | 0.57 | 0.46 | 0.03 | 0.87 |
| | H x EH x PL | 3.42 | 0.07 | 0.25 | 0.62 |
| Flower stem height | H | - | - | 1.03 | 0.32 |
| | EH | - | - | 1.94 | 0.17 |
| | PL | - | - | 4.19 | 0.05 |
| | H x EH | - | - | 0.21 | 0.65 |
| | H x PL | - | - | 0.75 | 0.40 |
| | EH x PL | - | - | 0.51 | 0.48 |
| | H x EH x PL | - | - | 0.00 | 1.00 |

#### Table S2

| Population | Ploidy | Evolutionary History | Region | GPS-N | GPS-E | Altitude | Characterization | Land use | Field site in Huber et al. 2016b |
|---|---|---|---|---|---|---|---|---|---|
| 1 | Diploid | allopatric | Switzerland - BE | 46.638 | 7.393 | 1031 | Rich pasture | Cut and autumn cow grazing | 18 |
| 2 | Diploid | Allopatric | Switzerland – BE | 46.632 | 7.386 | 935 | Intermediately rich pasture | Cut and autumn cow grazing | 20 |
| 3 | Diploid | allopatric | Switzerland - BE | 46.631 | 7.394 | 846 | Rich pasture | Cow grazing | 19 |
| 4 | Diploid | allopatric | Switzerland - BE | 47.069 | 7.424 | 539 | Rich pasture | Cow grazing & cutting | - |
| 5 | Diploid | allopatric | Switzerland - BE | 47.070 | 7.430 | 539 | Rich pasture | Cutting | - |
| 6 | Diploid | allopatric | Switzerland - BE | 47.072 | 7.439 | 539 | Rich pasture | Cutting | - |
| 7 | Diploid | sympatric | Switzerland - TG | 47.578 | 9.178 | 539 | Rich pasture | Cutting | - |
| 8 | Diploid | sympatric | Switzerland - TG | 47.580 | 9.161 | 569 | Rich pasture | Cutting | - |
| 9 | Diploid | sympatric | Switzerland - TG | 47.592 | 9.272 | 460 | Rich pasture | Cutting, occasional cow grazing | - |
| 10 | Diploid | sympatric | Switzerland - GB | 46.781 | 9.179 | 944 | Rich pasture | Cut and autumn cow grazing | 10 |
| 11 | Diploid | sympatric | Switzerland – GB | 46.759 | 9.205 | 854 | Relatively nutrient poor meadow | Cut | 13 |
| 12 | Diploid | sympatric | Switzerland - GB | 46.789 | 9.283 | 829 | Nutrient poor grassland | Cut | 12 |
| 13 | Triploid | sympatric | Germany - Spessart | 49.920 | 9.295 | 372 | Extensively used grassland | Cut and occasional sheep grazing | 1 |
| 14 | Triploid | sympatric | Germany – Spessart | 49.914 | 9.299 | 359 | Extensively used grassland | Horse grazing | 2 |
| 15 | Triploid | sympatric | Germany - Spessart | 49.910 | 9.299 | 335 | Intermediately rich pasture | NA | 3 |
| 16 | Triploid | allopatric | Germany - Würzburg | 49.737 | 9.604 | 345 | Extensively used grassland | Occasional grazing or cut | 8 |
| 17 | Triploid | allopatric | Germany – Würzburg | 49.726 | 9.651 | 323 | Extensively used grassland | Occasional grazing or cut | 9 |
| 18 | Triploid | allopatric | Germany - Würzburg | 49.721 | 9.731 | 283 | Extensively used grassland | Occasional grazing or cut | 7 |

**Figure S1**

a

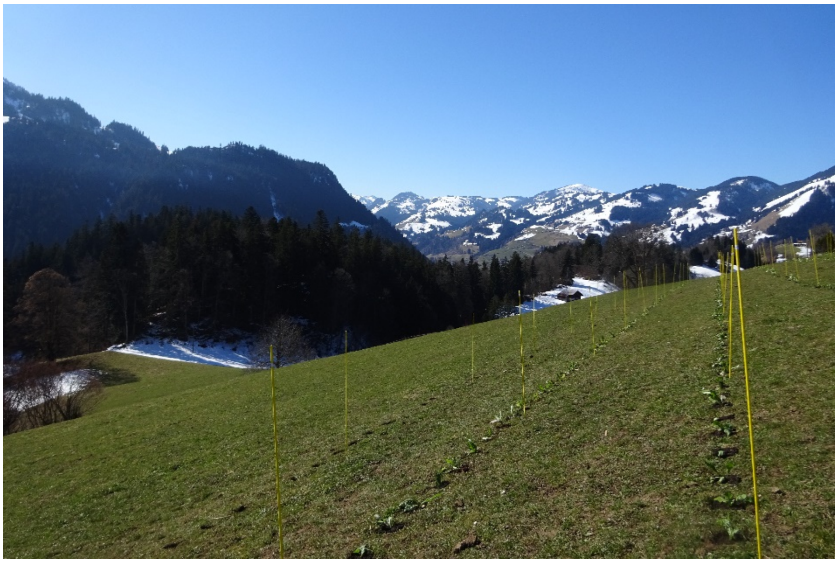

b

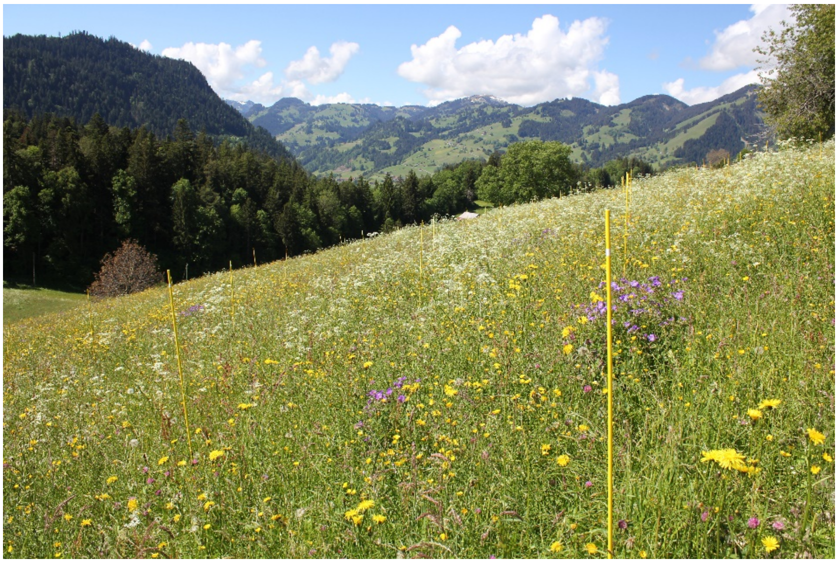

**Figure S2**

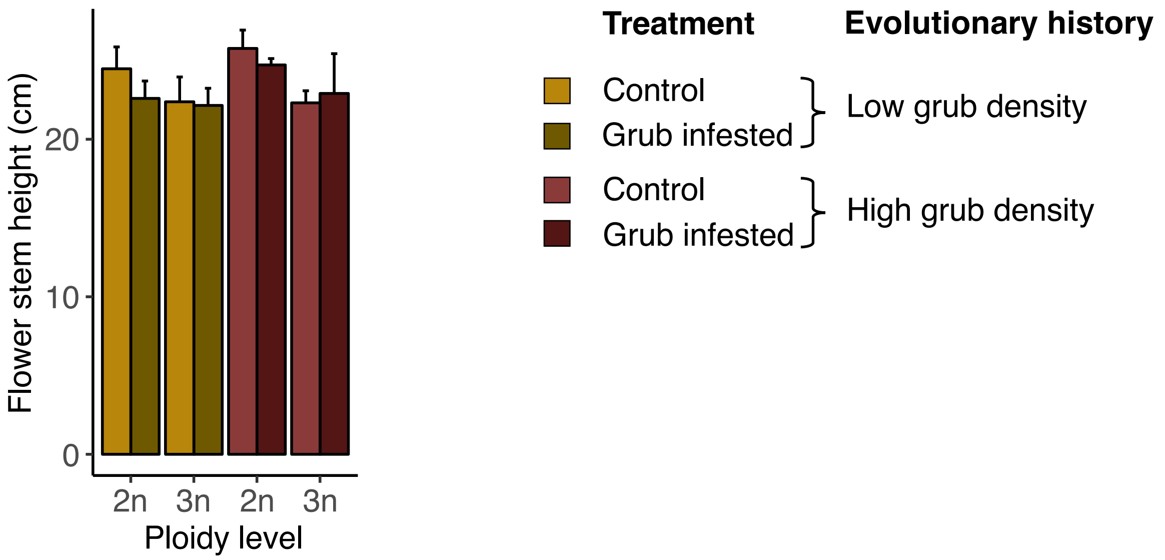
Flower stem height (cm)
0
10
20
2n
3n
2n
3n
Ploidy level
Treatment
Evolutionary history
Control
Grub infested
Low grub density
Control
Grub infested
High grub density

**Figure S3**

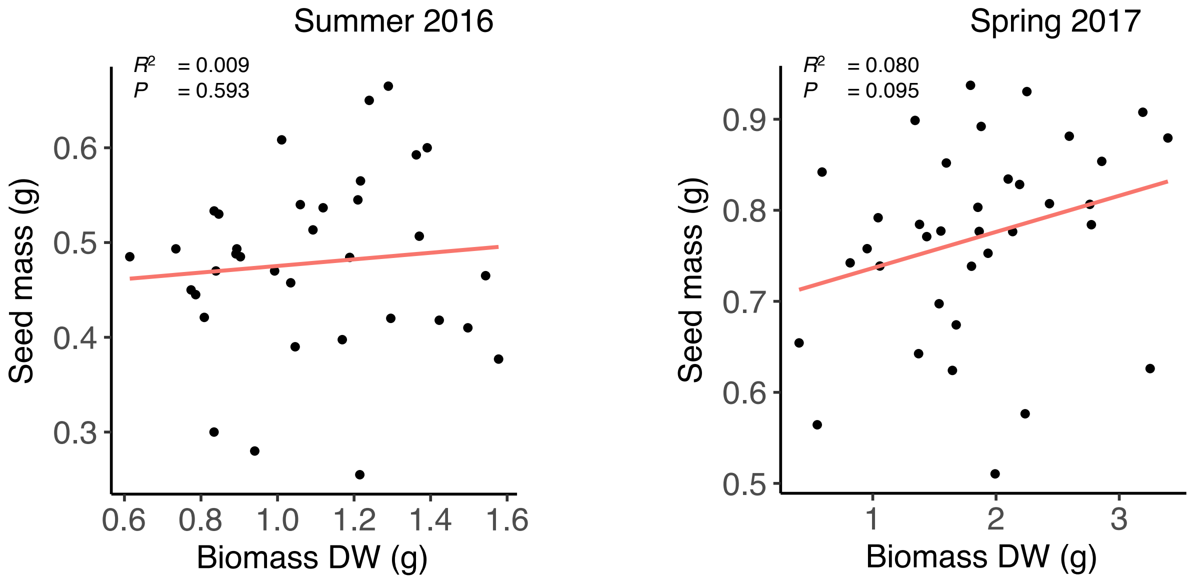

Summer 2016
$R^2$ = 0.009
$P$ = 0.593
Seed mass (g)
0.3
0.4
0.5
0.6
0.6
0.8
1.0
1.2
1.4
1.6
Biomass DW (g)
Spring 2017
$R^2$ = 0.080
$P$ = 0.095
Seed mass (g)
0.5
0.6
0.7
0.8
0.9
1
2
3
Biomass DW (g)
